# Robust assessment of the cortical encoding of word-level expectations using the temporal response function

**DOI:** 10.1101/2024.04.03.587931

**Authors:** Amirhossein Chalehchaleh, Martin Winchester, Giovanni M. Di Liberto

## Abstract

Speech comprehension involves detecting words and interpreting their meaning according to the preceding semantic context. This process is thought to be underpinned by a predictive neural system that uses that context to anticipate upcoming words. Recent work demonstrated that such a predictive process can be probed from neural signals recorded during ecologically-valid speech listening tasks by using linear lagged models, such as the temporal response function. This is typically done by extracting stimulus features, such as the estimated word-level surprise, and relate such features to the neural signal. While modern large language models (LLM) have led to a substantial leap forward on how word-level features and predictions are modelled, there has been little progress made towards the metrics used for evaluating how well a model is relating stimulus features and neural signals. In fact, previous studies relied on evaluation metrics that were designed for studying continuous univariate sound features, such as the sound envelope, without considering the different requirements of word-level features, which are discrete and sparse in nature. As a result, studies probing lexical prediction mechanisms in ecologically-valid experiments typically exhibit small effect-sizes, severely limiting the type of observations that can be drawn and leaving considerable uncertainty on how exactly our brains build lexical predictions. First, the present study discusses and quantifies these limitations on both simulated and actual electroencephalography signals capturing responses to a speech comprehension task. Second, we tackle the issue by introducing two assessment metrics for the neural encoding of lexical surprise that substantially improve the state-of-the-art. The new metrics were tested on both the simulated and actual electroencephalography datasets, demonstrating effect-sizes over 140% larger than those for the vanilla temporal response function evaluation.

## 1. Introduction

Speech comprehension requires our brains to transform sounds into meaning [1]. As part of that process, our brains must detect speech tokens such as words and interpret their meaning according to the prior context. In turn, that context can aid speech comprehension in challenging scenarios, such as noisy multi-talker environments [2]. The predictive processing theory [3, 4] offers a neurophysiological framework explaining how prior context might contribute to speech comprehension, proposing that sensory processing is underpinned by an active process involving the continuous attempts to predict upcoming sensory events [5, 6]. Strong evidence has been gathered indicating that this phenomenon extends to word predictions [7, 8], with stronger prediction errors leading to stronger neural activations measured with electroencephalography (EEG) and magnetoencephalography (MEG) [9, 10]. This relationship between neural activations and word prediction error, or lexical surprise, has been studied extensively by comparing the event-related potentials (ERP) in response to expected and unexpected words. As a result, a negative electrical deflection was measured at a post-stimulus latency of about 400 ms i.e., the N400 [7, 11].

The N400 has been studied widely by means of carefully crafted stimuli, for example impacting the contextual appropriateness (e.g., ‘I like my coffee with cream and sugar/socks’) [12]. Consequently, the N400 is typically estimated on unusual and short sentences, and by only considering the words targeted for the manipulation, typically the final word of each sentence, while ignoring all other words. Recently, methodologies were developed for modelling the relationship between continuous speech inputs and the neural signal as a linear time-invariant system. Such methods enable the study of word processing and prediction in ecologically-valid scenarios without the need for any manipulation of the speech material [5, 13, 14]. This estimate, called the Temporal Response Function (TRF), which was devised for studying the EEG/MEG encoding of the sound envelope [15], was only subsequently adopted to study linguistic encodings at various levels of abstraction, from phonology to semantics [5, 16-19].

Robust neural signatures of lexical surprise were measured via TRF estimations, exhibiting spatio-temporal patterns remarkably similar to those of the N400 ERP [12]. One marked distinction is, instead, that TRF estimations can capture subtle changes in lexical surprise that are naturally present in ecologically-valid speech, rather than relying on altered speech including artificially-placed surprising words. To reflect these remarkable similarities and distinctions with the N400 ERP, we refer to the TRF estimate of lexical surprise as the TRF-N400. The TRF approach is flexible in that it allows us to consider all words or subsets of interest, such as content words. Furthermore, it is important to note that, while we focus on lexical surprise for simplicity, our considerations on the TRF evaluation equally apply to similar features like word-level entropy and word dissimilarity.

The recent advances in large language models (LLM) have already contributed to the study of word-level processing, offering methods for the reliable and rapid estimation of word unexpectedness given the prior context [20, 21]. However, little progress has been made on the evaluation methods for word-level TRF, hampering our ability to probe the underlying neural processes and better understand how lexical predictions are actually built. First, the present study carries out a quantitative investigation of the weaknesses and limitations of existing TRF evaluation metrics for the study of lexical surprise, by using lagged ridge regression for deriving the TRFs via the mTRF-Toolbox [22]. That analysis pinpoints important limitations that substantially dampen the effect-sizes when probing lexical surprise with TRFs, which are due to key issues such as the assessment of lexical surprise encoding with suboptimal metrics, which were designed for continuous features like the sound envelope and are less appropriate for lexical surprise, which is discrete in time. Furthermore, the impact of collinearity is often ignored when building baseline models involving, for example, a random shuffling of the lexical surprise values. Based on these observations, we then introduce two novel evaluation metrics that are considerably more sensitive to lexical predictions. All analyses were carried out on a publicly available dataset recorded as participants listened to continuous speech. A simulated version of that dataset was also studied, where neural signals were constructed by combining an artificially-built neural response to speech and EEG noise, informing us on how accurately a TRF can retrieve the ground-truth neural response.

### 1.1 What evaluation metrics are typically adopted and their limitations

To determine if EEG/MEG signals encode lexical surprise or similar features (e.g., semantic dissimilarity), previous work using TRFs adopted multiple strategies. This section provides a brief introduction of common approaches and metrics, as well as an intuition of what limitations they might present. All the strategies covered in this study involve time-discrete features, consisting of vectors of zeros with the only non-zero values marking a linguistic event, such as word onsets. Those features can also be modulated by a second type of information, such as the word surprise. Hence, one challenge for TRF models using such time-discrete features is to disentangle the neural correlates of these two types of information.

The first approach consists of fitting a univariate TRF (uTRF), as done by Broderick and colleagues [13], and then observing the regression model weights to identify TRF components with distinctive spatio-temporal patterns that are consistent within and between listeners. This approach, which is also typical of ERP analyses, is complemented by an assessment of the neural signal variance explained by lexical surprise with that uTRF model, typically by measuring EEG/MEG prediction correlations with cross-validation. In that case, the lexical surprise uTRF is compared with a baseline model by looking at both model weights corresponding to lexical surprise and prediction correlations, where the baseline model is fit like the lexical surprise model, but after corrupting the lexical surprise information. That operation can be done by applying a random shuffling of the values, while preserving the word onset times, or by tampering with the LLM that generated the surprise values [21]. The second approach involves fitting a multivariate TRF (mTRF), as done by Di Liberto and colleagues [18] and [5], and the same metrics and baseline strategies mentioned above.

The intuition is that uTRFs are fit based on a single feature, such as lexical surprise, that is correlated with both acoustic and linguistic responses, meaning that the resulting models will likely capture both, hampering our ability to isolate neural correlates of lexical surprise. Using mTRF attempts to solve that issue, as lexical surprise is concatenated with nuisance features, such as word onset and the speech envelope, which are expected to absorb EEG/MEG variance that is unrelated to surprise. While this reasoning may sound intuitive and flawless, and despite its application in multiple studies from various research teams, there are at least two key issues with it. First, altering the lexical surprise values would likely lead to a change in the corresponding model weights. However, the weights for other features might also be affected due to collinearity. As a result, focusing only on the changes for the target feature while ignoring everything else might lead to incorrect conclusions.

The second key issue is that EEG/MEG prediction correlations are usually estimated by considering entire segments of data. While that is reasonable for envelope tracking studies, as the speech envelope is a continuous signal, word onset and lexical surprise are time-discrete vectors. For example, let’s consider two words *w1* and *w2* with a very large inter-word interval of *t12* = 3 second. In that case, large part of that EEG/MEG signal cannot be predicted by a lexical surprise feature, which would just inform us on the first few hundreds of milliseconds that follow *w1*. As a result, EEG/MEG prediction correlations would be calculated in a segment whose response is only partly affected by lexical surprise, substantially diluting the effect of interest. In other words, the intuition is that the prediction correlation metric is impacted by the word density, with slower speech leading to smaller effects of lexical surprise. The analyses that follow quantify and propose direct solutions to these issues.

## 2. Material and Methods

### 2.1 Natural-speech-EEG dataset

This study involves a re-analysis of a publicly available scalp EEG dataset, where neural signals were recorded as participants listened to narrative speech [13]. These data were part of a set of studies examining how human cortical signals encode acoustic and linguistic features of speech [16, 23, 24]. Nineteen participants (six female and thirteen male) aged between 19 and 38 years took part in the experiment. All participants were native English speakers, and reported normal hearing, normal or corrected-to-normal vision, and no history of neurological disorders. The experiment was conducted in a single session for each participant. EEG data were recorded as participants listened to an audiobook version of a popular mid-20th century American work of fiction (“The Old Man and the Sea”), read by a single professional male speaker. The audio stimuli were organized into 20 segments corresponding to the first chapters of the book, each with a duration of about 180 seconds. Segments were presented in a way that preserved the storyline, with neither repetitions nor discontinuities, and with an average speech rate of ∼210 words/min.

128-channel EEG data plus two mastoid channels were acquired at a rate of 512 Hz using a BioSemi ActiveTwo system. Triggers indicating the start of each trial were sent by the stimulus presentation computer and included in the EEG recordings to ensure synchronization. Testing was carried out in a dark, sound-attenuated room and participants were instructed to maintain visual fixation on a crosshair centered on the screen for the duration of each trial, and to minimize eye blinking and other motor activities. The present study utilized a version of the dataset that was shared according to the Continuous-event Neural Data structure (CND) [25, 26].

### 2.2 Simulated EEG data

One of the challenges of probing brain activity with technologies such as EEG is that the recorded neural signals are mixed with various sources of noise. Therefore, neural signatures derived by relating EEG and stimulus features likely reflect a combination of actual neural activity and EEG noise. Intuitively, a good assessment metrics would reflect how effective a model is at capturing the ground truth neural signal hidden behind the noise. However, since that ground truth signal is typically unavailable, we built a second dataset artificially with the CNSP simulation toolkit [27]. Specifically, the stimulus features used for generating the simulated EEG data were the Hilbert envelope of the speech sound and the lexical surprise vector, which was built using GPT-2 [20] (see **Section 2.5**). Stimulus features from the Natural-Speech-EEG dataset were convolved with predefined TRFs that were designed based on the literature. The simulated EEG data was derived by summing the two convolutions and, on top of that, EEG noise consisting of random segments of EEG data from the Natural-Speech-EEG dataset.

The predefined envelope and lexical surprise TRFs were generated with the following equations, approximating TRFs from previous studies [13, 28]:

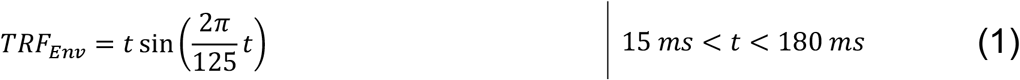

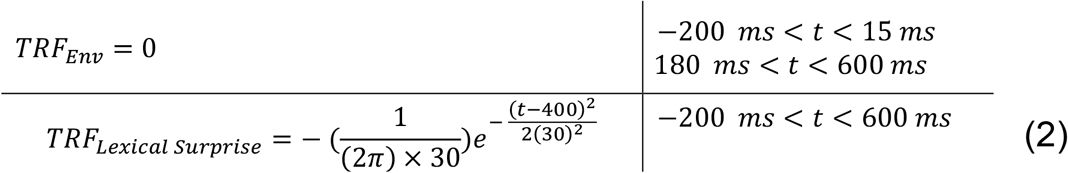

TRF_Env_ and the TRF_Lexical Surprise_ are shown in **figure 1(A)**. Please note that the outcome of our analyses on the simulated EEG dataset are insensitive to small changes in these artificial TRFs (e.g., time-shifting or scaling). Note that the resulting simulated EEG dataset has the same number of channels, trials, and participants as in the Natural-Speech-EEG dataset.

**Figure 1.**
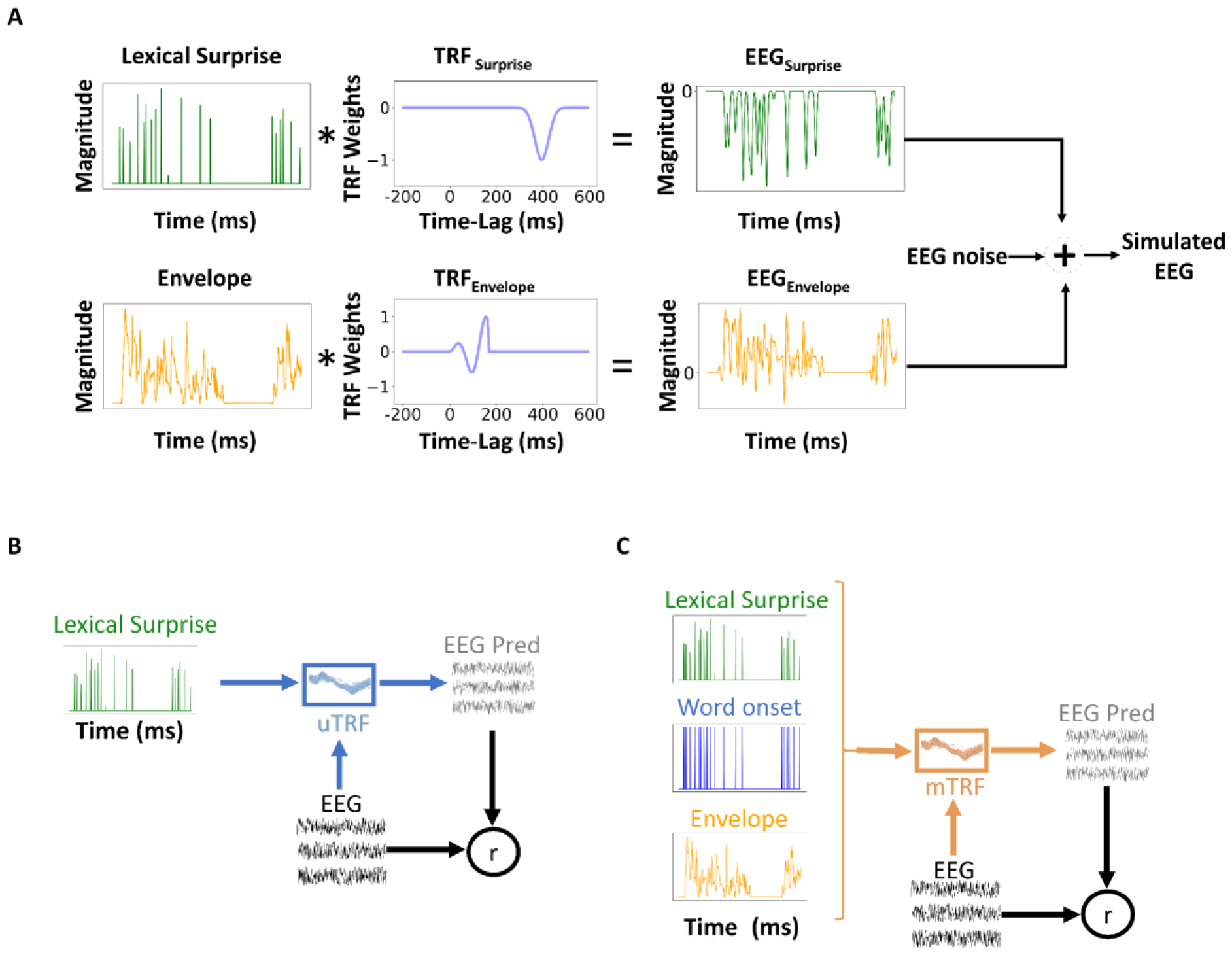
Methodological approach. **(A)** The simulated EEG dataset was generated by summing three signals. The first signal was the convolution of the speech envelopes from the Natural-Speech-EEG dataset [34] and a predefined impulse response, or temporal response function (TRF), with three main deflections representing the P1, N1, and P2 components of the TRF, approximating previous results on real data [13, 28]. The second signal was the convolution of lexical surprise with an artificially-built TRF with a single negative component at a latency of 400ms, broadly capturing previous results [35-37]. The third signal was EEG noise consisting of random segments of the EEG signal from the Natural-Speech-EEG dataset. **(B)** Forward univariate TRF models (uTRF) were fit to describe the mapping between lexical surprise and the EEG signal. The strength of the relationship between lexical surprise and EEG was assessed by comparing the lexical surprise uTRF with a baseline model, which was derived by fitting a second uTRF after randomly shuffling the lexical surprise values, while preserving their timing. EEG prediction correlations and TRF weights were used for the evaluation. **(C)** Forward multivariate TRF models (mTRF) were derived by relating a multivariate feature set consisting of the concatenation of envelope, word onset, and lexical surprise with the EEG signal. mTRFs were compared with a baseline model build by shuffling the lexical surprise values.

### 2.3 EEG preprocessing

EEG data were analyzed offline using MATLAB software (The MathWorks Inc.) according to the minimal preprocessing guidelines of the Cognition and Natural Sensory Processing initiative [29]. The same preprocessing pipeline and code were used for both datasets [25]. Signals were digitally filtered between 0.5 and 8 Hz using a Butterworth zero-phase filter, similar to previous studies [13, 30]. Both low- and high-pass filters had order 2 and were implemented with the function *filtfilt*, obtaining zero-shift phase filters. Signals were down-sampled to 128 Hz and re-referenced to the average of the mastoid channels. To identify channels with excessive noise, the time series were visually inspected, and the standard deviation of each channel was compared with that of the surrounding channels. Channels contaminated by excessive noise were recalculated by spline interpolating the surrounding clean channels in EEGLAB [31].

### 2.4 Temporal Response Function (TRF)

This study discusses evaluation metrics for TRF models, focusing on forward TRFs. A forward TRF can be described as a filter that linearly transforms a stimulus S(t) to a neural response R(t) over a specified series of time lags: R(t) = TRF_w_^*^S(t) + ε, where TRF_w_ are the weights of the filter at every time lag, and ε represents the residual of the prediction. Estimation of the TRF weights is done using regularized linear regression [22]. Previous studies have used this approach to investigate acoustic and linguistic processing with neural signals recorded as participants listened to continuous speech [5, 13, 30]. This approach can utilize a single stimulus feature at a time, or multiple such features simultaneously, leading to univariate TRF (uTRF) and multivariate TRF (mTRF) respectively. The latter has the advantage of availing of additional information for predicting the EEG data, which can lead to an improve ability of explaining EEG variance. Furthermore, mTRFs can more clearly inform on the distinct contributions to the EEG predictions of different features in cases of multicollinearity. Here, both the uTRF and mTRF approaches were explored to determine metrics that most reliably reflect the neural encoding of lexical surprise **figure 1(B, C)**.

The quality of the fit of TRF models is typically assessed with two types of metrics.

- The first metric is derived by building EEG predictions with cross-validation. The Pearson’s correlation of those predictions with the actual EEG data are then calculated on portions of signal that were not included in the model fit (test fold), producing correlation values for each EEG sensor, trial (e.g., chapter of an audio-book), and participant. For ease of analysis and visualization, prediction correlations are often averaged across EEG channels or calculated on selected scalp locations. This simplification comes at the cost of penalizing our assessment, in the former, and of limiting the analysis to a specific location while ignoring the others, in the latter.
- Ones it is verified that the TRF model explains some of the EEG variance by studying the EEG prediction correlations, it is then possible to study the weights of the regression model. The analysis of the TRF weights can inform us on which specific stimulus-EEG latencies and scalp areas are most relevant to their relationship (e.g.,∼400ms) [32]. This second metric is referred to as TRF weights.

### 2.5 Stimulus feature extraction

Three speech features were used to fit TRF models: speech *envelope, word onset, and lexical surprise*. The broadband amplitude *envelope* was computed by applying the Hilbert transform, capturing a key acoustic property of the speech material [33]. *Word onset* was defined as a vector containing ones and zeros, where ones denote the word onset. *Lexical surprise* serves as a proxy for semantic processing, as it quantifies how unexpected a word is depending on the proximal context. *Lexical surprise* obtained with LLM for all words in the speech has been shown to relate with the non-invasively recorded brain signal, enabling the isolation of word-level predictive processes from EEG [5]. Here, lexical surprise values were derived using GPT-2 [20], an open-source transformer-based LLM, which can be employed to gauge the surprise of each word based on its preceding context. The preceding context was built based on words heard within each particular chapter of the audio-book.

### 2.6 TRF model fit and evaluation

The neural encoding of word-level predictions can be studied with both uTRFs [30, 37, 38] and mTRFs [39-41]. To assess whether the EEG signals reflect lexical surprise, TRF results were compared when using lexical surprise vectors that did or did not match the speech material. Similar to previous TRF studies [42], mismatched surprise vectors were generated by randomly shuffling the order of the surprise values in the lexical surprise vectors, while preserving the onset times. To carry out statistical testing on individual participants, the shuffling and model-fit procedure was repeated 100 times, generating a null distribution for each participant. Note that this null distribution is stricter than simply mismatching the trial index for lexical surprise and EEG signal, as our procedure isolated the impact of the lexical surprise values, while everything else (i.e., word onset times and speech envelope) remained constant.

In uTRFs, the l*exical surprise* vectors and their shuffled versions were used to fit uTRFs. In that case, the two only differ in the order of the surprise values, while the timing was identical. Note that the shuffled lexical surprise vectors contain meaningful word onset timing, while the surprisal values can be seen as noise, as they are unrelated with the EEG signal. In mTRFs, models were fit by considering the three features (envelope, word onsets, and lexical surprise) simultaneously. In this case, speech envelope and word onsets acted as nuisance regressors, absorbing variance related to sound acoustics and word onsets. As such, the lexical surprise regressor was expected to more clearly capture variance that is unique to lexical surprise (and anything correlated to it that is not envelope and word onsets). Before analyzing the Natural-Speech-EEG dataset, uTRFs and mTRFs were evaluated on the simulated data, where we had full control on what information was and was not present in the EEG data (in this case, envelope and lexical surprise were encoded).

For simplifying the model evaluation, EEG prediction correlations calculated on single channels were averaged across all channels and trials, leading to a single value per participant. Regarding the model weights, averaging weights across EEG channels can be problematic instead, as positive and negative deflections in different scalp areas can cancel each other out. For that reason, our analyses of the TRF weights focused on a selected Centro-Parietal EEG channel, Pz, where the relationship between word-level predictions and EEG is known to be particularly strong [35, 37].

### 2.7 Two new metrics for evaluating word-level predictions

This study proposes two new evaluation metrics for assessing the neural encoding of word-level predictions. Note that these metrics could potentially be applied for the evaluation of TRFs for other common time-discrete features, for example at the level of phonemes [39, 43] or music notes [42, 44-46]. The first metric is specifically designed for improving the evaluation of surprise-like responses from the mTRF weights. The second metric aims to increase the sensitivity of EEG prediction correlations to temporally sparse events such as words, and it applies to both uTRF and mTRF methods.

#### ΔTime-constrained TRF weights, or ΔTC weights.

The first metric relies on TRF weights and is calculated on mTRF models. The observation is that Lexical surprise uTRF weights are not particularly sensitive to the surprise values (see **Figure 2A** and results section). The rationale is that the modulations relating to lexical surprise are likely small in natural speech, meaning that simple word-onset uTRFs capture the majority of the word responses already. To more clearly isolate the neural encoding of lexical surprise as opposed to word onsets, here we fit mTRF models by using lexical surprise and word onset features simultaneously. If the TRF model weights exhibit differences when examining the TRF weights for the two features, that would mean that the EEG signals encode correlates of lexical surprise. Otherwise, measuring undistinguishable TRF weights for lexical surprise and word onset would indicate that the specific lexical surprise is not encoded in the EEG signal. With this premise, the encoding of lexical surprise can be measured by subtracting the TRF weights for lexical surprise and word onset, and by calculating the absolute value of their difference at the latency where the major effect is expected, in this case 400ms. Specifically, weights were averaged in the window 350-450 ms to account for latency differences between participants. Note that our conclusions did not change for small changes in the selection of that window.

**Figure 2.**
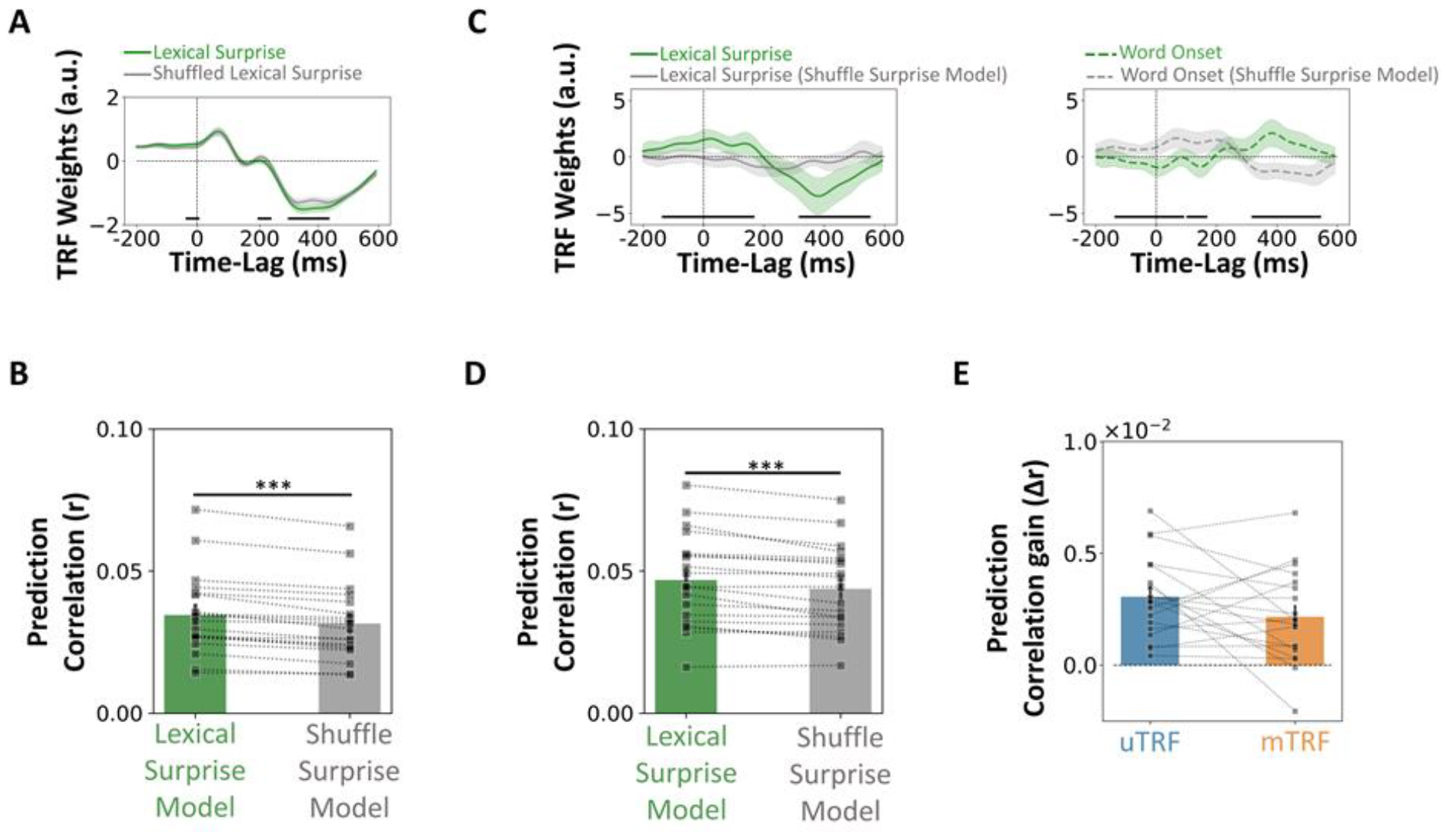
Probing the cortical encoding of lexical surprise with uTRF and mTRF on the Natural-Speech-EEG dataset. (**A**) uTRF weights (top) for lexical surprise were statistically significantly larger than for a shuffled version of lexical surprise (shuffle surprise model). The figures report the average TRF weights for individual features the EEG channel Pz across participants, with shaded areas indicating the standard error (SE). Black lines on the bottom of the panels indicate statistically significant differences between the TRF weights for the different features across time (Wilcoxon signed rank test, solid black line; p<0.05, FDR corrected). **(B)** EEG prediction correlations averaged across all EEG channels showed a statistically significant encoding of lexical surprise for uTRF models (bottom; ^***^p<0.001). Bars indicate the average across EEG participants and channels; dots refer to individual participants. **(C)** mTRF weights of lexical surprise and shuffle surprise model. (left) lexical surprise weights, (right) word onset weights. Statistically significant effects of lexical surprise also emerged for mTRFs when comparing TRF weights of lexical surprises and word onset in the lexical surprise model and the shuffle surprise model (solid black line; p <0.05, FDR corrected). Colors indicate the mTRF model i.e., green for the lexical surprise model, grey for the shuffle surprise model. **(D)** EEG prediction correlations averaged across all EEG channels showed a statistically significant encoding of lexical surprise for mTRF models (^***^p<0.001). **(E)** No statistically significant differences emerged when comparing EEG prediction correlation gains (i.e., the increase when using lexical surprise values rather than shuffled values) between uTRF and mTRF models.

#### Time-constrained (TC) EEG prediction correlation.

The second metric relies on EEG prediction correlations. Feature vectors capturing word-level information are typically sparse as they code the information of interest, such as lexical surprise, time-locked to the word onsets. Considering a word rate between about 100 and 260 words per minute across different speakers, speech categories and languages [47], and considering that TRF-N400 evoked component can extend approximately between 200 and 600 ms [13, 30, 41]. Under these general assumptions, the worst-case scenario would be that only 20 seconds every minute of EEG data would actually reflect lexical surprise. As such, 66% percent of the datapoints used for calculating the EEG prediction correlation would not capture the effect of interest, substantially diluting that effect. Here, we propose to calculate the EEG prediction correlations by only considering the datapoint that might reflect the target effect. This simple modified metric is applied at the evaluation stage, and it does not affect the TRF model fit, meaning that it can be used to evaluate both uTRF and mTRF models. The code used for deriving this metric has been shared on the GitHub of the mTRF-Toolbox (function *mTRFcrossval_multimetric*).

### 2.7. Statistical analysis

Statistical significance in group-level analysis was assessed through pair-wise Wilcoxon signed-rank tests applied to both EEG prediction correlations and mTRF weights In the single-subject level analysis (**figure 4(D) and S2(D)**), we conducted 100 permutations for each participant, systematically shuffling the values of lexical surprise in each iteration. This process generated a null distribution for each subject, allowing us to calculate statistical significance at the individual level. Wilcoxon signed rank tests were used for pair-wised comparisons. Correction for multiple comparisons was applied where necessary via the false discovery rate (FDR) approach. A two-way repeated measures ANOVA was used to assess the effects of within and between factors. The values reported use the convention *F*(*df*_*numerator*_, *df*_*denominator*_). FDR-corrected Wilcoxon tests were used after ANOVA for post hoc comparisons.

## 3. Results

### 3.1 The challenge of probing lexical surprise in the human cortex with univariate TRFs

The relationship between lexical surprise vectors and EEG was evaluated with uTRFs on both simulated and real EEG data. For the simulated EEG data, TRF weights were compared between the lexical surprise model and the null (shuffled lexical surprise) model. Wilcoxon signed-rank test showed that there is a negative deflection for the lexical surprise model around 400ms (**figure S1(A)**; Wilcoxon signed-rank test on the average TRF weights in the 300-500ms lag window: *p* < 0.001, *d* = 0.4 FDR-corrected Wilcoxon tests were carried out on individual lags with *p* < 0.05). Prediction correlation comparison between the lexical model and the shuffle surprise model were greater for the lexical model (**figure S1(B)**; Wilcoxon signed-rank *p* < 0.001, *d* = 9.72).

Similar outcomes emerged when analysing real EEG data. TRF weights at channel *Pz* were compared between lexical surprise and the shuffled surprise models, showing a stronger negative deflection for the lexical surprise model at latencies close to 400ms (**figure 2(A)**; Wilcoxon signed-rank test on the average TRF weights in the 300-500ms lag window *p* < 0.001, *d* = 0.22; FDR-corrected Wilcoxon tests were carried out on individual lags with *p* < 0.05). EEG prediction correlations were also larger for the lexical surprise model than the shuffled surprise model, leading to a statistically significant difference (**figure 2(B)**; *p* < 0.001, *d* = 0.17).

These results are in line with the literature in that a statistically significant encoding of lexical surprise is measured. As in previous work, both this result corresponded with small effect sizes (d ⪅ 0.2) for both TRF weights and EEG prediction correlations metrics. The analysis in **Section 3.3** aims at deriving evaluation metrics that are more sensitive to lexical surprise, leading to larger effect sizes.

### 3.2 Probing lexical surprise with multivariate TRFs

Further analyses were carried out to evaluate whether the flexibility of multivariate TRFs can improve the evaluation of lexical surprise encoding in EEG signals. One of the advantages of mTRFs is that, by using multiple stimulus features at once, the different contributors to the prediction can be more clearly separated [15, 18]. Here, lexical surprise mTRFs were derived by including envelope, word onset, and lexical surprise features as the input, while the shuffle surprise model utilized a shuffled version of the lexical surprise vectors. The intuition is that lexical surprise and word onset feature vectors would capture different EEG variance only when the surprise values are meaningful and encoded in the EEG signal. This intuition was tested by comparing the TRF weights averaged between 300 and 500ms on the simulated and channel *Pz* for the real EEG data across the two *features* (surprise and word onset) and two *models* (lexical surprise and shuffled surprise.

Results on the simulated EEG data showed main effects of feature, model, and their interaction (repeated measures ANOVA, feature: *F*(1,18)=0.50, *p*=0.48; model: *F*(1,18)=4.81, *p* < 0.05; feature^*^model: *F*(1,18)=68.73, *p* < 0.001, with statistically significant differences between lexical surprise and word onset (*p* < 0.001, *d* = 3.50) and between shuffled surprise and word onset (*p* < 0.001, *d* = 2.57).

For the real EEG dataset, we found main effects of feature, model, and their interaction (repeated measures ANOVA, feature: *F*(1,18)=20.68, *p* < 0.001; model: *F*(1,18)=5.22, ^*^*p* < 0.05; feature^*^model: *F*(1,18)=22.98, *p* < 0.001, with statistically significant differences between lexical surprise and word onset (*p* < 0.001, *d* = 2.57) but not between shuffled surprise and word onset in the shuffle surprise model (*p* = 0.104, *d* = 0.49). TRF weights corresponding to lexical surprise in the lexical surprise and shuffle surprise model showed statistically significant differences at individual lags (**figure 2(C-left)**; FDR-corrected Wilcoxon signed rank test, *p* < 0.05). Also, TRF weights related to word onset in lexical surprise and shuffle surprise showed statistically significant differences at individual lags (**figure 2(C-right)**; FDR-corrected Wilcoxon signed rank test, *p* < 0.05).

EEG prediction correlations for the simulated EEG dataset were larger for the lexical surprise model than the shuffle surprise model (**figure S1 (D)**; Wilcoxon signed rank test; *p* < 0.001, *d*= 3.64). EEG prediction correlations were also compared between the uTRF and mTRF models showing, as expected, larger prediction correlations for the mTRF model (*p* < 0.001, *d* = 275.6). While this difference captures the effect of using three speech features, envelope, word onset and lexical surprise, simultaneously rather than only lexical surprise, we also quantified the EEG variance explained by lexical surprise by subtracting the EEG prediction correlations for the lexical surprise and shuffle surprise models, where lexical surprise information was present and absent respectively. This EEG prediction gain showed statistically significant difference between uTRFs and mTRFs, with uTRFs having larger prediction gain (**figure S1 (E)**; *p* < 0.001, *d* =12.67).

EEG prediction correlations for the real EEG dataset were also larger for the lexical surprise model than the shuffle surprise model (**figure 2(D)**; Wilcoxon signed rank test; *p* < 0.001, *d* = 0.18). EEG prediction correlations as expected showed larger prediction correlations for the mTRF model compared to the uTRF model (*p* < 0.001, *d* =0.83). While this difference captures the effect of using three speech features simultaneously rather than only lexical surprise, the EEG prediction gain did not show any statistically significant difference between uTRFs and mTRFs (**figure 2(E)**; *p* = 0.332, *d* = 0.249).

### 3.3 Isolating robust neural metrics of lexical surprise

The previous sections indicate that TRF metrics described in the literature (i.e., EEG prediction correlations and TRF weights) can be used to probe the lexical surprise generated by the human cortex during speech comprehension. However, comparing lexical surprise and shuffle surprise models exhibited small effect-sizes, when using such metrics. To magnify our ability to measure lexical prediction processes, two novel TRF metrics are introduced.

The **ΔTC weights** metric, which is calculated on the weights of the mTRF model, showed larger values for the lexical surprise model than the shuffle surprise model (**figure S2(A);** Wilcoxon signed-rank test: *p* < 0.001, *d* = 3.13) on the simulated EEG dataset. The same result also emerged on the Natural-Speech-EEG dataset (**figure 3(A)**, *p* < 0.001, *d* = 2.09). While this effect was only evaluated on TRF weights around the 400ms time-latency, where the impact of lexical surprise was expected to be strongest, the contrasts of TRF weights for lexical surprise and word onsets is reported in **figure 3(C)** across all the time-latencies in the TRF models. That visualization further highlights the value of studying that contrast, which is shows statistical significant effects for the lexical surprise model but not for the shuffled lexical surprise model (FDR-corrected Wilcoxon signed rank test, *p* < 0.05).The ΔTC weights metric was also sensitive to lexical surprise at the level of individual participants, with 16 out of 19 of them exhibiting larger values for the lexical surprise model than the shuffle surprise model (**figure 3(D-left)**, FDR-corrected Wilcoxon signed-rank test, *p* < 0.05); whereas the same analysis on the univariate model results showed statistically significant effects in only 9 out of 19 participants (**figure 3(D-right)**, *p* < 0.05).

**Figure 4.**
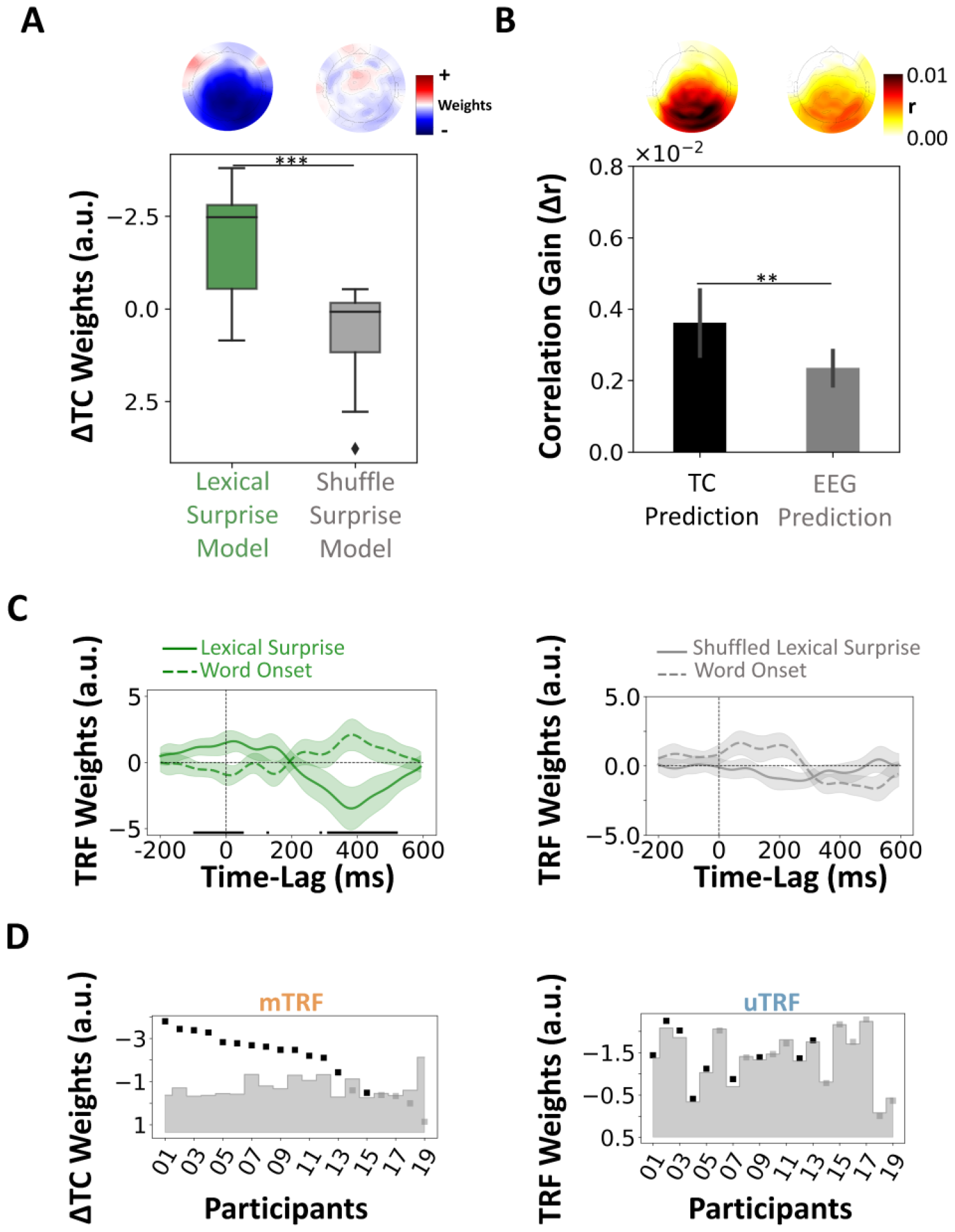
Robust assessment of the EEG encoding of lexical surprise. **(A)** Evaluation of the ΔTC weights metric on the Natural-Speech-EEG dataset. The box plot shows the distribution of ΔTC weights for the lexical surprise model and the shuffle surprise model (^***^p < 0.001) (y-axis inverted). The distribution of the ΔTC weights across the scalp sensors are shown above the box plot. **(B)** TC and EEG prediction correlation gains (lexical surprise vs. shuffled surprise model), when using mTRFs. The bar plots show the mean correlation gains (± SE) across participants, EEG channels and trials when using each of the metrics (^**^p < 0.01). Topographies of the TC and EEG prediction correlation gains are shown above the bar plots. **(C)** Pz mTRF weights for the lexical surprise model (left) and shuffle surprise model (right). Black lines on the bottom of the plots indicate statistically significant differences between the TRF weights for lexical surprise and word surprise (p < 0.05, FDR corrected). Green and grey colors indicate the mTRF lexical surprise and shuffle surprise models respectively. **(D)** Individual participant level results for the ΔTC weights in an mTRF analysis (left) and the TRF weights in a uTRF analysis (right). All weights were calculated for the P_z_ EEG channel here. ΔTC weights were obtained by considering the window-size of 300-500ms. The upper limit of the shaded grey area shows the 95^th^percentile of the null distribution obtained for individual participants. Black data-points are reported for statistically significant results (FDR corrected, p < 0.05).

The second novel metric, **TC prediction correlation**, consists of identifying time-points unrelated with the target effect by design, and then excluding those time-points when calculating the EEG prediction correlation. This time-constrained correlation metric led to a substantial enhancement of the EEG prediction correlation gain values on both the simulated EEG dataset (**figure S2(B)**, Wilcoxon signed-rank test, *p* < 0.001, *d* = 2.05) and the natural speech EEG dataset (**figure 3(B)**, *p* = 0.009, *d* = 0.44). Correlation gain results showed a 144% increase in effect size when using TC correlation metric for model evaluation.

## 4. Discussion

This study identified limitations with the use of TRFs with time-discrete stimulus features, such as lexical surprise. We then proposed two new metrics that tackle those issues directly. The new metrics were tested on both simulated and actual EEG data, exhibiting effect-sizes that were over 100% larger than those for the vanilla TRF evaluation. The first metric magnifies the effect on the mTRF weights by contrasting weights for word onsets and lexical surprise. The intuition lexical surprise vectors capture word onsets and surprise information. So, if the surprise values were meaningless (e.g., if the values were shuffled), similar TRF weights would emerge for lexical surprise and word onsets. Meaningful surprise values would instead lead to a different set of weights for the two features, which is why we expected their contrast to be representative of lexical surprise encoding. Effect sizes computed with this metric demonstrated significantly greater magnitude (d=2.09) in comparison to univariate model evaluations utilizing the TRF weights (d=0.22). The second metric, TC prediction correlation, improves the EEG prediction correlation metric by accounting for the temporal sparsity of the word onsets and, specifically, by only considering time-points that can actually be influenced by lexical surprise. Effect sizes derived from this metric also exhibited a considerably larger magnitude compared to mTRF model assessments utilizing prediction correlation for evaluation, marking a 144% increase in effect size.

One of the key challenges when measuring linguistic level processing with EEG is that a large portion of the EEG response is explained by the acoustic changes in the sound. Lexical surprise has the advantage of producing a neurophysiological component, the TRF-N400, that is clearly distinct from the typical envelope TRF, both in terms of temporal and spatial patterns, making it possible to separate the two with mTRF models. Instead, it is less clear how effective the TRF approaches discussed here would be at determining how exactly the surprises are built. For example, it is possible to build different hypotheses by using distinct LLMs, or by altering the amount of context available for the prediction, similarly to previous music neurophysiology research with Markov chains [28] and music transformers [46]. The new metrics in the present study increase the sensitivity to lexical surprise, making that type of fine-grained comparisons more feasible. Therefore, we expect future work to explore this direction and to provide valuable insights on how context is built and used during speech processing. Recent developments have already shed some light on that question, leading to the promising result that the internal organisation of the rapidly advancing LLMs is getting progressively closer to the speech processing pathways in the human cortex [21]. The assessment metrics proposed in the present study are expected to contribute to that line of work with a different angle into that question, shedding light on how linguistic context is built and then used to process speech.

The results of this study can be summarized into recommendations for future research. The first observation is that the literature is quite inconsistent in the way the TRF-N400 is evaluated, challenging the comparison and aggregation of different studies. Some studies consider word onsets and their modulation together [13], while others attempt to separate two neural signatures by relying on different approaches for calculating a baseline. In our view, the random shuffling baseline presented here, which was already used by other previous studies, could serve as a consistent baseline across different studies, as the shuffling procedure could equally applied to any modulated time-discrete feature. Therefore, we encourage the use of this baseline in the future. Indeed, multiple baselines can be calculated and should be considered, depending on the goals of each study. For example, it has been suggested that corrupting the model (e.g., LLM) in some ways [21] (e.g., retraining the model with random data, reducing the available context) might be a more conservative baseline than shuffling, as the latter would completely destroy the any regularity in the temporal structure. Nonetheless, that baseline and its effectiveness would depend on the specific language model and the goals of the evaluation. One final recommendation based on our results is that measuring how the lexical surprise weights are affected by a baseline, like a shuffled surprise, is insufficient and potentially deceiving in case of strong collinearities in the feature-set. The weights of other features, in fact, would also likely be affected, as measured in **figure 3(B)**. Therefore, we recommend observing the entirety of the change in the regression weights when considering such baselines, for example by adopting the procedure proposed in **figure 4(A)**.

## Acknowledgements

We thank Franklenin Sierra and Aoife Igoe for their help with the code for generating the lexical surprise

