## Supplementary Figures for "Robust assessment of the cortical encoding of word-level expectations using the temporal response function"

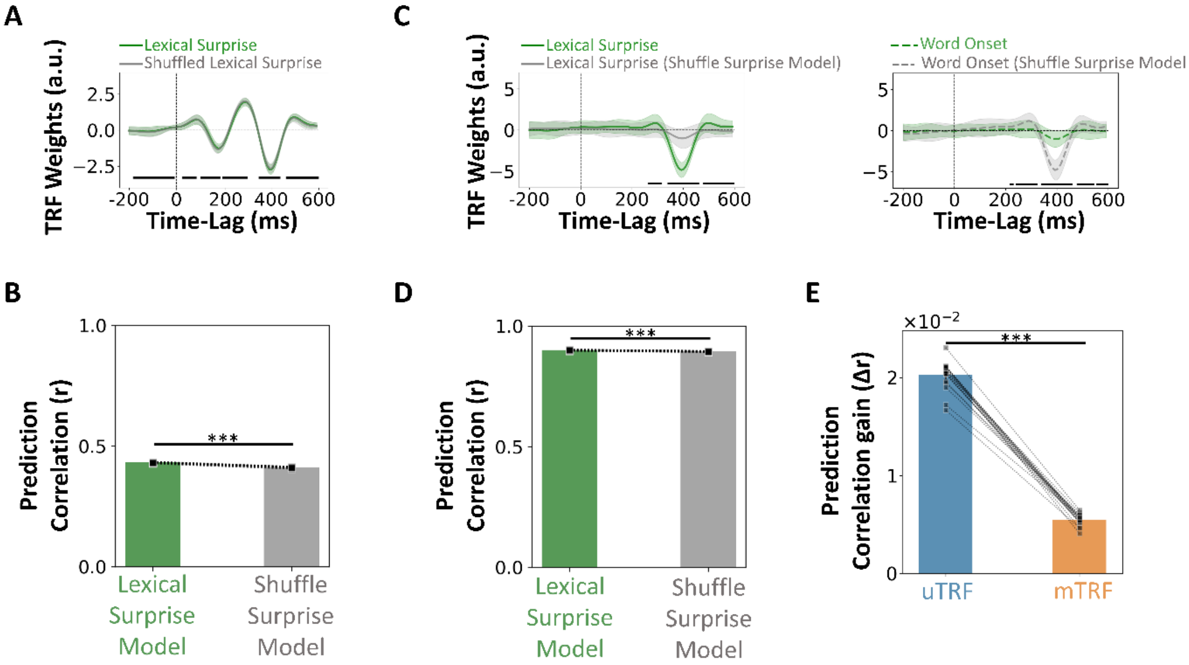


***Figure S1. Probing the cortical encoding of lexical surprise with uTRF and mTRF on simulated EEG dataset.*** *(**A**) uTRF weights (top) for lexical surprise were statistically significantly larger than for a shuffled version of lexical surprise (shuffle surprise model). The figures report the average TRF weights for individual features the EEG channel* $P_z$ *across participants, with shaded areas indicating the standard error (SE). Black lines on the bottom of the panels indicate statistically significant differences between the TRF weights for the different features across time (Wilcoxon signed rank test,* $p < 0.05$*, FDR corrected). (**B**) EEG prediction correlations for simulated EEG channel showed a statistically significant encoding of lexical surprise for uTRF models (bottom;* $^{***}p < 0.001$*). Bars indicate the average across EEG participants; dots refer to individual participants. (**C**) mTRF weights of lexical surprise and shuffle surprise model. (left) lexical surprise weights, (right) word onset weights. Statistically significant effects of lexical surprise also emerged for mTRFs when comparing TRF weights of lexical surprises and word onset in the lexical surprise model and the shuffle surprise model (*$p < 0.05$*, FDR corrected; black lines on the bottom of the plots indicate statistically significance). Colors indicate the mTRF model i.e., green for the lexical surprise model, grey for the shuffle surprise model. (**D**) EEG prediction correlations for simulated EEG channel showed a statistically significant encoding of lexical surprise for mTRF models (*$^{***}p < 0.001$*). (**E**) statistically significant differences emerged when comparing EEG prediction correlation gains (i.e., the increase when using lexical surprise values rather than shuffled values) between uTRF and mTRF models (*$^{***}p < 0.001$*).*

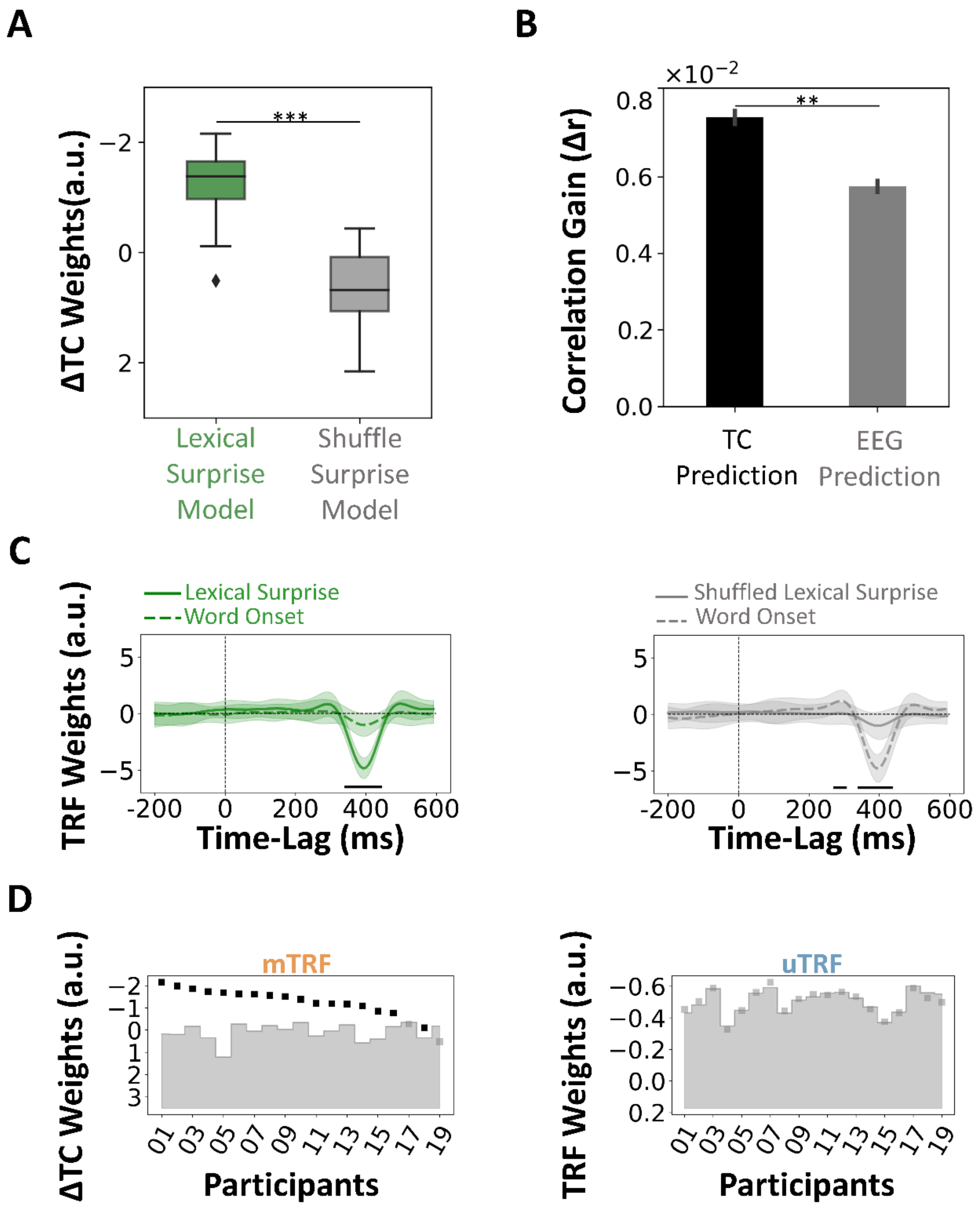


***Figure. S2. Robust assessment of lexical prediction processes on simulated EEG dataset. (A)*** *Performance of the ΔTC weights metric. The box plots show the distribution of ΔTC weights for the semantic model and the shuffle surprise model (*$^{***}p < 0.001$*) (y-axis inverted). Topography images for the ΔTC weights are shown alongside the related boxplots.* ***(B)*** *Performance of the TC prediction correlation and the EEG prediction correlation metrics in term of correlation gain, when using an mTRF approach. The bar plots show the mean correlation gain (± SE) across participants, electrodes and trials when using each of the metrics (*$^{**}p < 0.01$*). Topography images for the correlation gains in TS prediction correlation and the EEG prediction correlation metrics are shown alongside the related bar plots.* ***(C)*** *mTRF weights for the lexical surprise model (right) and shuffle surprise model (left). Statistically significant effects of lexical surprise also emerged for mTRFs when comparing TRF weights for lexical surprise and word onset (*$p < 0.05$*, FDR corrected; black lines on the bottom of the plots indicate statistically significance). This difference emerged in the lexical surprise and in the shuffle surprise model with different directions. Colors indicate the mTRF model i.e., green for the lexical surprise model, grey for the shuffle surprise model.* ***(D)*** *Individual participant level results for the ΔTC weights in an mTRF analysis (left) and the TRF weights in a uTRF analysis (right), for the* $P_z$ *electrode and window sizes 200ms centered around 400ms. The upper limit of the shaded grey area shows the* $95^{th}$ *percentile for individual*

*participants obtained from the null distributions (chance level). Grey squares show insignificant participants after correcting for multiple comparison (FDR corrected, $p < 0.05$).*
